# Development of a Novel Benzodiazepine to Delineate Peripheral GABA-A Signaling Mechanisms in Visceral Pain Syndromes

**DOI:** 10.1101/2025.06.08.658501

**Authors:** Michael S. Poslusney, Qian Li, Ingrid P. Buchler, Yifang Huang, Liansheng Liu, Yaohui Zhu, Subhash Kulkarni, Gregory Carr, Adrienne DeBrosse, Noelle White, Diane Peters, James C. Barrow, Pankaj J. Pasricha

## Abstract

**Background and Aims:** Visceral pain is a cardinal symptom of many disorders affecting the gut. Modulators of gamma-aminobutyric acid (GABA) such as benzodiazepines may attenuate colonic pain but the specific contribution of peripheral GABAA receptors remains unclear as these agents have prominent central effects.

**Methods:** Using medicinal chemistry optimization of the benzodiazepine scaffold, we developed a novel and potent benzodiazepine-based positive allosteric modulator (PAM) of GABAA receptors, Li633, with no significant central nervous system (CNS) penetration.

**Results:** The locomotor activity of rats placed in an open field was unchanged with Li633 at doses up to 30 mg/kg, confirming its lack of a CNS effect. LI-633 produced robust potentiation of GABA-induced inward current, with EC50 values ranging from 8 nM (α5β2γ2) to 128 nM (α3β2γ2). In vitro electrophysiological studies confirmed its ability to reduce excitability of human dorsal root ganglion (DRG) neurons. LI-633 potentiated muscimol-induced GABAergic currents in rat DRG neurons in a dose-dependent manner, with an EC_50_ of 70.4 nM. In vivo, LI-633 significantly attenuated visceral hypersensitivity and pain behavior in a rat model of irritable bowel syndrome (IBS) and functional dyspepsia (FD). In the IBS model, administration of the drug also resulted in decreased excitability of colon-specific DRG neurons and significantly reduced the colonic afferent response to balloon distention as measured by recordings of neural activity in dorsal ganglia rootlets.

**Conclusions:** These findings highlight the potential of targeting peripheral GABAA receptors for pain management in IBS and other disorders associated with visceral hypersensitivity.

## Introduction

Chronic pain involves complex interactions between the peripheral and central nervous systems.^1^ In the periphery, tissue damage or inflammation can lead to sensitization of nociceptors, making them more responsive to stimuli. This peripheral sensitization can trigger changes in the central nervous system (CNS), particularly in the spinal cord and brain, leading to central sensitization. It has been difficult to assess the relative contributions of these two nervous systems to chronic pain as almost all analgesics cross the blood brain barrier and therefore can modulate the function of both. However, such insight has important therapeutic implications-targeting the predominant component (peripheral or central) and avoiding off-target adverse effects or targeting both and providing more effective relief.^2^

This is particularly relevant in the development of drugs for disorders characterized by chronic visceral pain, such as irritable bowel syndrome (IBS) and functional dyspepsia which together affect more than 10% of the population in the USA and world-wide.^3^ Uniquely, gastrointestinal sensory afferents originate and terminate in two large, complex nervous systems (the enteric nervous system or ENS, and the CNS, respectively) with large overlaps in both neurotransmitters and their receptors, including, amongst others, serotonin, dopamine and gamma-aminobutyric acid (GABA).^4^ GABA is the endogenous agonist for two distinct families of receptors, designated GABA_A_ and GABA_B_. GABA_A_ receptors comprise a large group of ligand-gated channels that allow the passage of chloride ions when open. These heteropentameric channels are generally composed of two alpha subunits, two beta subunits, and a single gamma subunit.^5^ GABA is widely distributed in the CNS where it exerts a hyperpolarizing, inhibitory effect contributing to the well-known sedating effects of drugs such as benzodiazepines and other positive allosteric modulators (PAMs) of the GABA_A_ receptor. In the periphery, GABA_A_ signaling can be either hyperpolarizing or depolarizing, depending on the local chloride gradient across the membrane of the target cell.^6^ Although its function in the gastrointestinal tract has yet to be fully worked out, the ENS has the highest concentration of GABA amongst all peripheral organs and there is intriguing evidence for a role for GABA signaling both as part of IBS pathophysiology and as a potential therapeutic target.^7, 8^

However, the site of action of both the clinical and pre-clinical drugs are difficult to assess because the modulators used (benzodiazepines and muscimol, respectively) cross the blood brain barrier. Although the potential utility GABA_A_-PAMs for the treatment of chronic pain is supported by a large body of literature that shows that these compounds exhibit anti-allodynic and anti-hyperalgesic effects,^9, 10^ some of these effects are likely to also be mediated centrally by α_2_, and to a lesser extent α_3_-containing, GABA_A_ receptors within the spinal cord.^11^ Nevertheless, recent evidence supports a prominent role for peripheral GABA_A_ receptors in mediating analgesia that appears to be distinct from their central effects. Dorsal root ganglia (DRG) and associated glial cells have been demonstrated to express functional GABA_A_ receptors as well as all of the requisite “molecular machinery” needed to synthesize, store, and release GABA; the exogenous delivery of GABA, or GABA agonists to the DRG has been shown to effectively attenuate nociceptive behavior in models of inflammatory and neuropathic pain.^12^ With respect to visceral sensitization, muscimol has been shown to inhibit attenuate colonic afferent activity in isolated colon-pelvic nerve preparations and behavioral pain responses to noxious colonic distention in normal mice.^13^ Further, both diazepam and GABA attenuate visceral hypersensitivity in mice with acute colitis^14^.

We therefore hypothesized that enhancement of GABAergic signaling only within peripheral primary sensory neurons, and without engagement of central receptors, could be an effective strategy to treat visceral pain associated with irritable bowel syndrome. Herein, we report the characterization of a promising lead GABA-A PAM, LI-633, that is pharmacologically similar to classical benzodiazepines, but with highly restricted access to the CNS. This agent reduces excitability in human sensory neurons in vitro and is effective in preclinical models of aspects of irritable bowel syndrome. The development of this compound has the potential to identify more precisely the respective contributions of the gut and the brain to the pathogenesis of chronic symptoms such as pain as well as spur the development of novel therapies for the same.

## Materials and Methods

### Rat pharmacokinetic experiments

Experiments were carried out at Pharmaron (Beijing, China) using male Sprague-Dawley rats (see Supplementary Materials for details).

### Nonspecific GABA Binding Assay

Tritiated flunitrazepam displacement from rat brain tissue was performed as described by Speth et al,^15^ at CEREP, Celle l’Evescault, France.

### Manual patch-clamp in recombinant cell lines

Experiments were carried out by B’Sys GmbH (Witterswil, Switzerland). CHO cells stably expressing human GABA_A_ α_1_ß_2_γ_2_ receptors and LTK cells stably expressing human GABA_A_ α_2_ß_2_γ_2_, α_3_ß_2_γ_2_, or α_5_ß_2_γ_2_ receptors were used for all experiments (see Supplementary Materials for details).

### Off-target binding panel

LI-633 was tested against a panel of 87 enzymes, receptors, and ion channels (Eurofins Panlabs, Taipei, Taiwan). LI-633 was tested in duplicate at a concentration of 10 uM.

### Rat dorsal root ganglia (DRG) cell electrophysiology

Experiments were carried out at Metrion Biosciences (Cambridge, UK). Dorsal root ganglion neurons were isolated from P8-P12 Sprague-Dawley rats, were prepared and maintained in culture for up to four days on glass coverslips. Whole-cell manual patch clamp recordings were obtained from cell bodies (see Supplementary Materials for details).

### Human DRG Electrical Field Stimulation Study

Experiments were carried out at AnaBios (San Diego, USA). Tissue preparation: Human DRGs obtained from consenting donors were transferred into a dissection vessel containing a cold (4°C), fresh proprietary dissection solution. DRGs were maintained completely submerged in dissection solution and dissected appropriately. Each DRG was enzymatically dissociated as per AnaBios’ proprietary methodologies (see Supplementary Materials for details).

### Open-field Locomotor Activity

Eight-week-old male rats were ordered from Charles River (Wilmington, MA, USA) and allowed to acclimate to our animal facility for one week. Rats were then handled daily (∼1 min) for 7 consecutive days prior to behavioral testing. On the test day, the rats were transported to the behavioral testing room, weighed (290-330g), and left to acclimate for one hour. After the one-hour waiting period, the rats were dosed (PO; volume: 5 mL/kg) with either vehicle (5% DMSO in phosphate-buffered saline; pH = 8) or a single dose of LI-633 (3, 10, or 30 mg/kg) and then placed into a novel, acrylic open-field arena (445 mm length X 445 mm width) for two hours. The sessions were recorded and locomotor activity was analyzed using Topscan (CleverSys, Inc., Reston, VA, USA) automated behavioral analysis software.

### Neonatal Irritable Bowel Syndrome (IBS) Model and Pseudoaffective Responses to Noxious Colorectal Distention

IBS rat model was generated as we have previously described.^16^ Briefly, male Sprague Dawley rat pups at postnatal day 6 were purchased with dams (Harlan Laboratories). At postnatal day 10-12, pups received a colorectal infusion of 0.2 ml of 0.5% acetic acid (AA) and were allowed to grow up until 8-12 weeks of age. For acute treatment, adult IBS or control rats were treated with LI-633 (1, 3, 10 and 30 mg/kg, 5 ml/kg, PO) or vehicle (5% DMSO in 0.1 M Na-phosphate buffer (pH 8). As a positive control, another group of IBS rats was treated with buprenorphine (0.5 mg/kg, subcutaneously). One hour later, hyperalgesia was assessed by visceromotor reflex (VMR) responses (measured by electromyographic recordings of the external oblique abdominal muscle) to colorectal distention (CRD). For repeated treatment, control and IBS rats were treated with LI-633 (1, 3, 10 and 30 mg/kg, 5 ml/kg, PO) or vehicle (5% DMSO in 0.1 M Na-Phosphate buffer (pH 8). As control, another group of IBS rats was treated with buprenorphine (0.5 mg/kg, subcutaneously) once a day for 5 days. The VMR response to CRD was then measured on day 6, 24 hours after the last administration.

### Ex-vivo measure of excitability on colonic sensory neurons in IBS rats

To label DRG neurons that innovated with distal colon, a retrograde dye, DiI ((1,1’-dioleyl-3,3,3’,3-tetramethylindocarbocyanine methanesulfonate, 10 mg / ml methanol) was injected into distal colon wall (1µl / site x 10 sites) of rats, previously sensitized with saline or AA as neonates as described previously by us.^17^ Two weeks later, rats were treated with vehicle or LI-633 (1 or 10 mg/kg, 5 ml/kg, PO) once a day for 5 days. On the day after the last treatment, rats were administrated with one dose of the treatment and were sacrificed one hour later. DRGs (L4-S3) were harvested, dissociated and plated, and whole-cell voltage patch-clamp recordings of the cultured DRG neurons were conducted in DiI-positive neurons as previously described by us.^18^

### Dorsal rootlet single nerve unit recording

To determine the effect of LI633 on peripheral sensory signal to the central nervous system, we tested the activity of dorsal rootlet (DR) single fibers using single-unit afferent recordings as previously described by us.^19^ Briefly, after anesthesia, the T13-S1 spine was exposed, mounted on a stereotaxic frame and fixed by clamping. Then, the spinal cord L1-6 were exported by the laminectomy. In an oil pool, the dorsal roots (DR) were then exposed by carefully cutting and deflecting the dura mater using fine forceps. To record the afferent fibers activity of the dorsal rootlets, the lumbar 5 DR was first cut centrally as close to the cord entry as possible and freed from the spinal cord. The nerve activity was recorded at the distal ends using a silver hook electrode. Single units that innervate the colon were identified by consistent spike rate in response to CRD. Signals were amplified with Iso-DAM8A Bio-amplifier (WPI) and analyzed using SPIK 2 software program (Cambridge Electronic Design, UK). A 20 second baseline was recorded followed by CRD for 20 seconds. Two pressures, 40 mmHg and 80 mmHg, were used for CRD. The single unit nerve activity was measured before and 10, 30, 60 and 90 minutes after administration of LI633 (5mg/kg, iv). The data were normalized to the baseline before administration.

### Effects of LI633 treatment visceral hypersensitivity in a rat model of functional dyspepsia (FD)

A rat model of FD was generated using transient neonatal gastric irritation previously described by us.^19, 20^. Ten-day-old male rats received 0.2 mL 0.1% iodoacetamide (IA) in 2% sucrose daily by oral gavages for 6 days; controls received 2% sucrose. Adult FD and control rats (∼ 12 Weeks old) were administered with LI633 (10 mg/kg, PO) or vehicle (5% DMSO in 0.1 M Na-phosphate buffer, pH 8). One hour after the treatment, the hyperalgesia of the rats was examined by the visceral motor reflex (VMR) responses to gastric distention (GD), as measured by electromyographic recordings of the acromotrapezius muscle.

### Statistical analysis

Data are expressed as mean ± SEM of the group (*n =* 6–8, unless otherwise noted). Data were analyzed by Student t-Test, 1-way or 2-way ANOVA using SigmaPlot (Systat software Inc.), unless otherwise specified. If a significant difference was detected, a Student-Newman-Keuls post hoc test was used to evaluate differences between individual groups. For all tests, *P <* 0.05 was considered significant.

## Results

### Lead Discovery characterization

Lead compound LI-633 (Figure 1A) emerged from medicinal chemistry optimization of the benzodiazepine scaffold for GABA_A_ positive allosteric modulation potency, efficacy, pharmacokinetics, and minimal brain penetration. Examination of the cryo-EM structure of the GABA-A receptor with bound diazepam indicates that the lactam moiety of diazepam is solvent-exposed.^21^ In several other prototypical benzodiazepines such as midazolam and rilmazolam, this lactam is replaced by a fused 5-membered heterocycle. Recognizing that this solvent-exposed portion of the molecule would tolerate many substituents, we sought to introduce polar or charged functional groups at this position. The carboxylic-acid substituted imidazole present in LI-633 was ultimately found to have an acceptable balance potency and peripheral restriction and was advanced into further characterization. In the ^3^H-flunitrazepam displacement assay, LI-633 was found to be a potent binder of GABA_A_ (Figure 1B), with a K_i_ of 21 nM. This value is comparable to the affinity of diazepam, which has a K_i_ of 7.1 nM.

**Figure 1.**
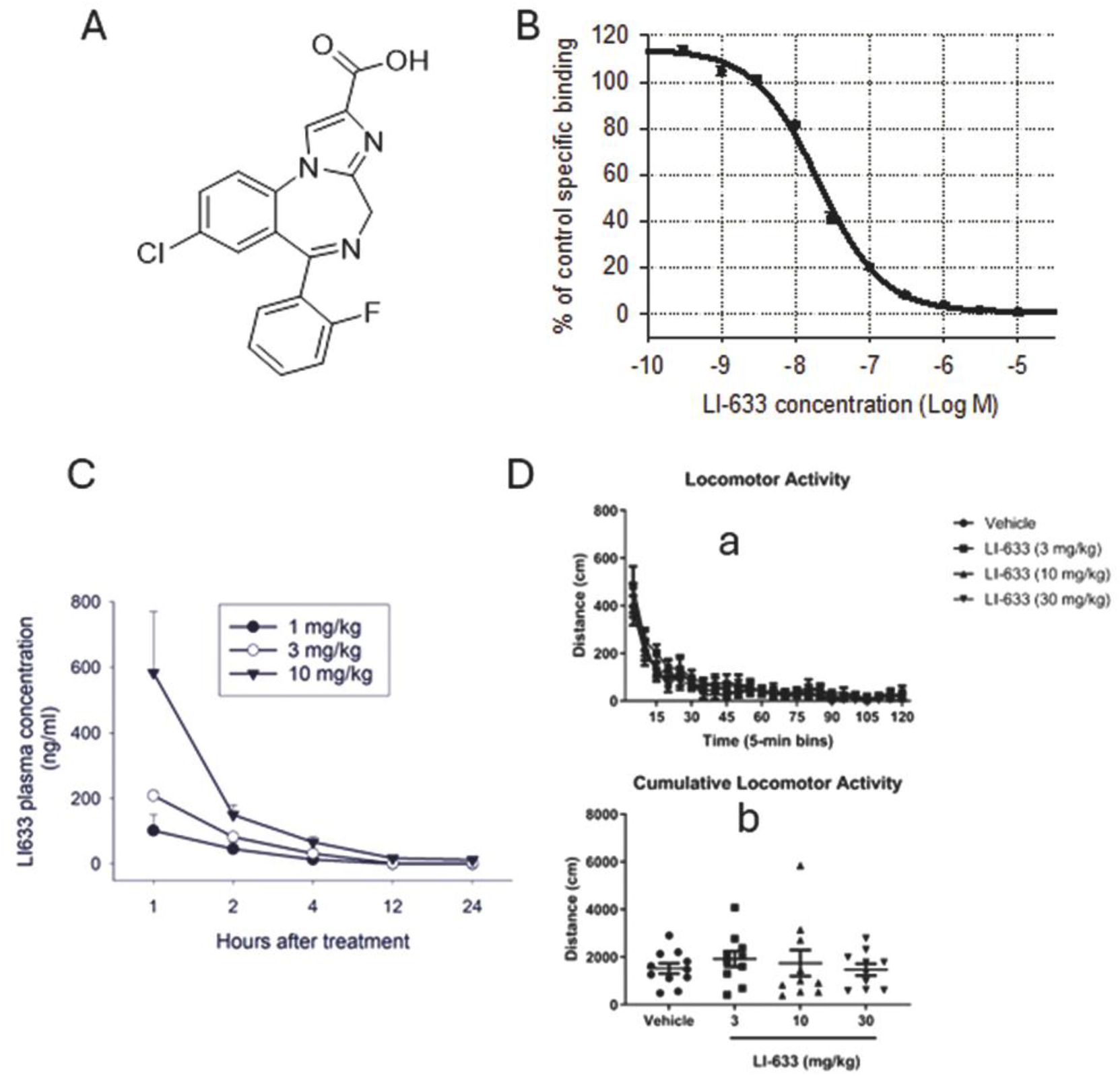
(**A**) Structure of LI-633; (**B**) Displacement of ^3H^-flunitrazepam from rat brain tissue by LI-633; **(C**) Time-course of plasma concentrations of LI633 in Sprague-Dawley rats following single oral doses (1, 3, 10 mg/kg); and (**D)** Locomotor activity of rats following administration of LI-633 (a) Locomotor activity measured in 5-min bins over 120 minutes (b) Cumulative locomotor activity during the 120-minute observation period. n = 11, 10, 10, 10 for vehicle, 3 mg/kg, 10 mg/kg, and 30 mg/kg LI-633 groups respectively. Data are presented as mean ± SEM.

To determine its functional effect across various GABA_A_R subunits, and to reveal any selectivity for particular subtypes, LI-633 was tested by manual patch clamp electrophysiology in cell lines expressing α_1_β_2_γ_2_, α_2_β_2_γ_2_, α_3_β_2_γ_2_, and α_5_β_2_γ_2_, receptors. As shown in Table 1, LI-633 produced robust potentiation of GABA-induced inward current, with EC_50_ values ranging from 8 nM (α_5_β_2_γ_2_) to 128 nM (α_3_β_2_γ_2_).

**Table 1:**
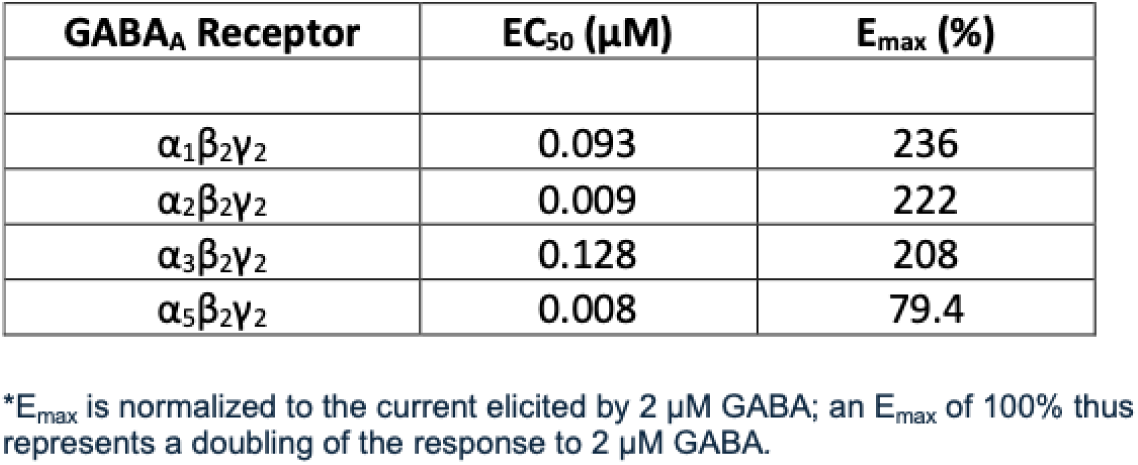
Activity of LI-633 against different GABA_A_ receptor subtypes.

| **Table 1:** Activity of LI-633 against different GABA<sub>A</sub> receptor subtypes |  |  |
| --- | --- | --- |
| **GABA<sub>A</sub> Receptor** | **EC<sub>50</sub> (μM)** | **E<sub>max</sub> (%)** |
| $\alpha_1\beta_2\gamma_2$ | 0.093 | 236 |
| $\alpha_2\beta_2\gamma_2$ | 0.009 | 222 |
| $\alpha_3\beta_2\gamma_2$ | 0.128 | 208 |
| $\alpha_5\beta_2\gamma_2$ | 0.008 | 79.4 |

\*E<sub>max</sub> is normalized to the current elicited by 2 μM GABA; an E<sub>max</sub> of 100% thus represents a doubling of the response to 2 μM GABA.

LI-633’s specificity was tested using a panel of 87 diverse targets, including enzymes, GPCRs, and transporters in rat brain preparations. As expected, LI-633 resulted in displacement of the benzodiazepine antagonist flumazenil and benzodiazepine flunitrazepam. No other significant interactions were observed in this assay, including GABA-B receptors (data not shown).

The *in vitro* ADME (absorption, distribution, metabolism, and excretion) profile of LI-633 is shown in Table S1 (Supplementary materials). Apparent permeability was assessed in MDR1-MDCK cell monolayers and was found to be low. The efflux ratio in MDR1-MDCK cells was 2.8, suggesting modest efflux by the P-gp transporter. LI-633 was unchanged after incubation with human or rat hepatocytes or liver microsomes. In a panel of 5 CYP450 enzymes, LI-633 displayed no inhibition up to a concentration of 50 uM. Moderate plasma protein binding was observed in human (10.9% unbound) and rat (18.6% unbound).

### Pharmacokinetic profile

The plasma levels of LI-633 were measured after administering oral doses of 1, 3, or 10 mg/kg in Sprague-Dawley rats (Figure 1C). LI-633 was rapidly absorbed, reaching a maximum plasma concentration less than 1 hour after administration.

Additionally, the concentration of LI-633 within tissue of the small and large intestine, dorsal root ganglion (DRG), and sciatic nerve (SN) was measured at three time points following an oral dose of 10 mg/kg. The LI633 concentrations were modest in DRG and sciatic nerve (the ratio of DRG/plasma and SN/plasma were 0.367 and 0.283 respectively one hour after administration while not detectable after 4 and 12 hours). However, a large amount of LI-633 was retained in the small and large intestines. The ratio of small intestinal to plasma concentration was 49, 58.1 and 41.7 at 1, 4 and 12 hours after administration respectively, whereas the ratio of colon to plasma was 0.916, 124 and 374 at 1,4 and 12 hours respectively after administration.

The potential for CNS penetration of LI-633 was assessed following a single 100 mg/kg oral dose. The total brain/plasma ratio at 1-hour post-dose was 0.021 (K_p)_, and the unbound brain/plasma ratio was 0.015 (K_pu,u_). The CSF/plasma ratio 1-hour post-dose was determined to be 0.007 (K_p_). These results demonstrate that LI-633 does not readily cross the blood-brain barrier, even after a high oral dose.

### Open-field locomotor activity

Following single oral doses of 3, 10 or 30 mg/kg, the locomotor activity of rats placed in an open field was unchanged compared to vehicle-treated animals, as shown in Figure 1D, indicating that LI-633 does not produce measurable sedation at these doses.

### Electrophysiological effects of LI-633 on rat DRG neurons

Next, we evaluated the ability of LI-633 to potentiate muscimol-induced currents in rat DRG neurons. Representative current traces are shown in Figure 2Ai. In a concentration-response experiment, the GABA_A_ selective agonist muscimol was found to induce GABA_A_R activation with an EC_50_ of 5.7 uM (Figure S1), consistent with previous reports.^22, 23^ In the presence of an EC_30_ concentration of muscimol (3 uM), LI-633 potentiated GABAergic currents in a dose-dependent manner, with an EC_50_ of 70.4 nM and an E_max_ value of approximately 100% (Figure 2Aii). Thus, LI-633 acts as a potent GABA_A_ PAM in native rodent tissue, in agreement with in vitro EC_50_ results from heterologous expression systems (Table 1).

**Figure 2.**
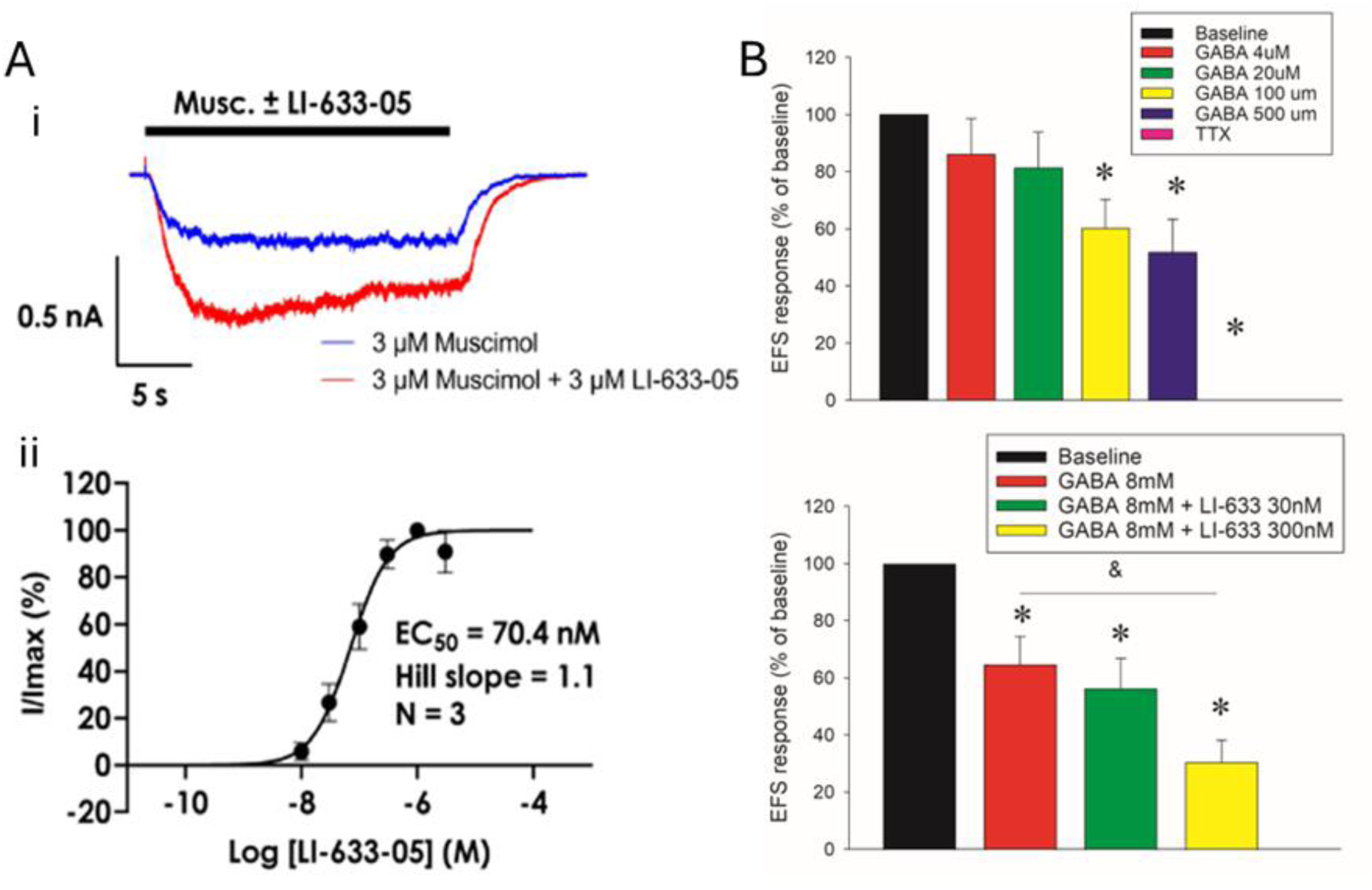
(**A**) LI-633 potentiates muscimol-induced inhibition of current in DRG neurons. **(a)** Representative recording from a rat DRG neuron, showing potentiation of muscimol-induced current by LI-633; **(b)** Concentration-dependence of potentiation of currents by LI-633 in rat DRG neurons. (**B**) LI633 potentiated GABA-induced reduction in EFS-induced excitability in human DRG neurons. **(a)**. GABA dose-dependently decreased excitability of human DRG neurons in response to EFS; **(b).** LI-633 potentiates the effect of submaximal dose of GABA on human DRG neuronal excitability in response to EFS. Data are presented as mean ± SEM of % of baseline of EFS response. *: Significant difference from baseline, P<0.05; &: Significant difference between two bars P<0.05. TTX = tetrodotoxin.

### Effects of LI-633 on excitability of human DRG neurons

Electrical field stimulation (EFS) is an established technique for studying neuronal excitability, and has been adapted for use with human DRG neurons.^24^ Using stimulation with voltage of 1500-2000 mV, among the sub-population of neurons that were GABA-responsive, excitability (as measured by calcium fluorescence using Fluo-8) was decreased by GABA in a concentration-dependent manner (Figure 2B). As positive control, TTX (Tetrodotoxin), a Na channel blocker, completely diminished the EFS response in the DRGs. At a submaximal concentration of GABA (8 uM), the effect of GABA was significantly potentiated by 300 nM, but not by 30 nM LI-633 (Figure 2B). Time-matched vehicle control experiments indicated that rundown from repeated electrical stimulation accounted for less than 25% of the reduction in EFS response. In all experiments, the positive control tetrodotoxin suppressed the EFS response to near zero.

### Visceromotor response to noxious colorectal distension in a rat model of IBS

The effects of LI-633 on hyperalgesia were tested in a well-established and accepted rat model of IBS-like pain.^16^ To assess acute drug treatment effects, IBS and control rats were treated with 0, 1, 3, 10, or 30 mg/kg of LI-633 (5 ml/kg, PO). Another group of IBS rats was treated with buprenorphine (0.5 mg/kg, SC), an opioid analgesic, as a positive control. One hour after the treatment, colorectal pain sensitivity was assessed by measuring the visceromotor reflex (VMR) response to colorectal distention (CRD). The CRDs were conducted with 4 pressures, 20, 40, 60 and 80 mmHg. The VMR responses to CRD were measured by EMG. In each treatment group, the EMG responses for 4 pressures were calculated into area under the curve (AUC) to present the data for each treatment. As shown in Figure 3A (left), IBS rats showed a significant increase in VMR response to CRD. Buprenorphine completely blocked the VMR response to CRD in IBS rats. Treatment with LI633 dose-dependently reduced the hyperalgesia in the IBS rats. Comparison of the AUC between groups, One-Way ANOVA revealed a significant difference between groups (F_(6,47)_=18.472, P<0.001). The post-hoc test showed that treatment with 3, 10 and 30 mg/kg LI633 significantly reduced hyperalgesia in IBS rats.

**Figure 3.**
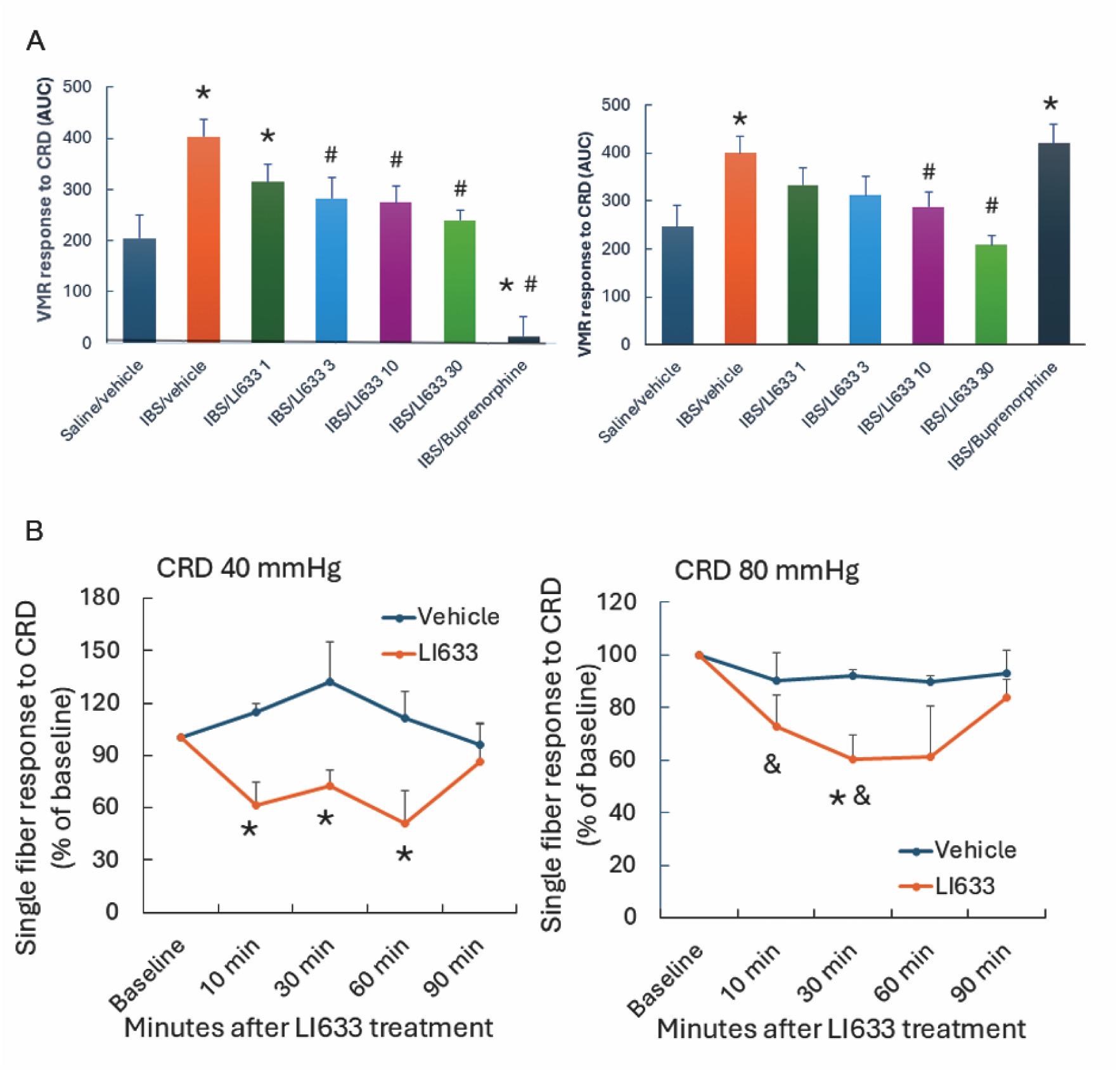
(A) Acute (left) and repeated (right) treatments of LI633 dose-dependently reduce the hyperalgesia in rat IBS model as assessed by visceral motor reflex (VMR) responses to colorectal extension (CRD) with 20, 40, 60 and 80 mmHg pressures. Data are expressed as an area under the curve (AUC) of EMG response to four pressures in each treatment group. *: Significantly different from Saline/Vehicle group, P< 0.05 by Student Newman-Keuls test. #: Significantly different from AA/Vehicle, P< 0.05 by Student Newman-Keuls test. (**B**) LI-633 reduces activity of dorsal root single fiber in response to CRD with 40 mmHg (left) and 80 mmHg (right) in IBS rats. Data are presented as single fiber activity in response of CRD relative to before (baseline) administration of vehicle or LI-633 (5 mg/kg, i.v). *: Significantly different from vehicle treatment at same time point, P< 0.05 by Student Newman-Keuls test.

To determine the effects of repeated treatment of LI-633, we administered LI-633 at the same doses as in the acute study for 5 days, and VMR responses to CRD were tested on the day after the last treatment. As shown in Figure 3A (right), five-day treatment of LI-633 dose-dependently reduced VMR-response to CRD in IBS rats. One-Way ANOVA reveals a significant difference between groups (F_(6,46)_ = 5.034, P<0.001). The IBS rats were significantly increased VMR response to CRD. Treatment with LI-633 at all doses diminished the significant increase in VMR response relative to saline/vehicle group, whereas treatment with 30 mg/kg had significantly lower CRD responses than that in IBS-vehicle group (P<0.001 by Student Newman-Keuls test). Interestingly, the buprenorphine treated IBS rats showed a rebound effect on the day after the last treatment, with visceral hyperalgesia exceeding that of the IBS-vehicle control rats. These studies demonstrate that LI-633 reduces hyperalgesia in a model of IBS, suggesting that it may be a good candidate for treatment of visceral pain.

### Dorsal rootlet responses to LI-633 in a rat model of IBS

To determine whether the effects of Li-633 on reduction of visceral hyperalgesia in IBS model is mediated by dorsal root-spinal pathway, we measured the activity of dorsal rootlet single nerve fibers in response to colorectal distention (CRD). Colorectal-innervated single fibers in the dorsal root nerve were identified by their response to CRD. The nerve activity in response to CRD was recorded before and after systemic administration of vehicle or LI-633 in IBS rat model (Figure 3B). Administration of vehicle did not induce significant changes in the nerve activity in response CRD, whereas LI-633 (5mg/kg, iv) significantly reduced the activity of dorsal rootlet single nerve fibers in response to CRD at both 40 mmHg (Two-Way ANOVA: Main effect of time: F_(4,25)_ = 0.706 P=0.595, Main effect of treatment F_(1,25)_ = 15.593, P < 0.001 n=4) and 80 mmHg (Two-Way ANOVA: Main effect of time: F_(4,25)_ = 1.964, P=0.131, Main effect of treatment F_(1,25)_ = 7.138, P = 0.013 n=4).

Thus, these results indicate that LI-633-induced reduction of hyperalgesia in IBS rats is associated with attenuation of peripheral nociceptive signals.

### Effects of LI-633 on excitability on colonic sensory neurons in the rat IBS model

To further establish the peripheral site of action of LI-633, we performed whole-voltage patch clamping on colon-specific spinal sensory neurons isolated from the DRG of IBS and control rats treated with 1 and 10 mg/kg of LI-633 (See Methods). Although resting membrane potential was significantly less negative in AA-sensitized rats, LI-633 did not have any effect (Two-way ANOVA: main effect of model F_(1,119)_ = 15.28, P<0.001, main effect of treatment: F_(2,119)_ = 0.23, P=0.79, Figure 4). On the other hand, rheobase (the amount of current needed to elicit an action potential spike) was significantly lower in AA-sensitized rats and reversed by treatment with LI-633 (Two-way ANOVA: main effect of model F_(1,119)_ = 20.54, P<0.001, main effect of treatment: F_(2,119)_ = 4.17, P=0.018, Figure 4). Further, the number of spikes induced by 2x-rheobase current was significantly higher in AA-sensitized rats relative to saline rats and also decreased by treatment with Li633 (Two-way ANOVA: main effect of model F_(1,119)_ = 22.22, P<0.001, main effect of treatment: F_(2,119)_ = 5.83, P=0.004, Figure 4). These results indicate that Li633 attenuates the increased excitability of colonic sensory neurons in this IBS model.

**Figure 4.**
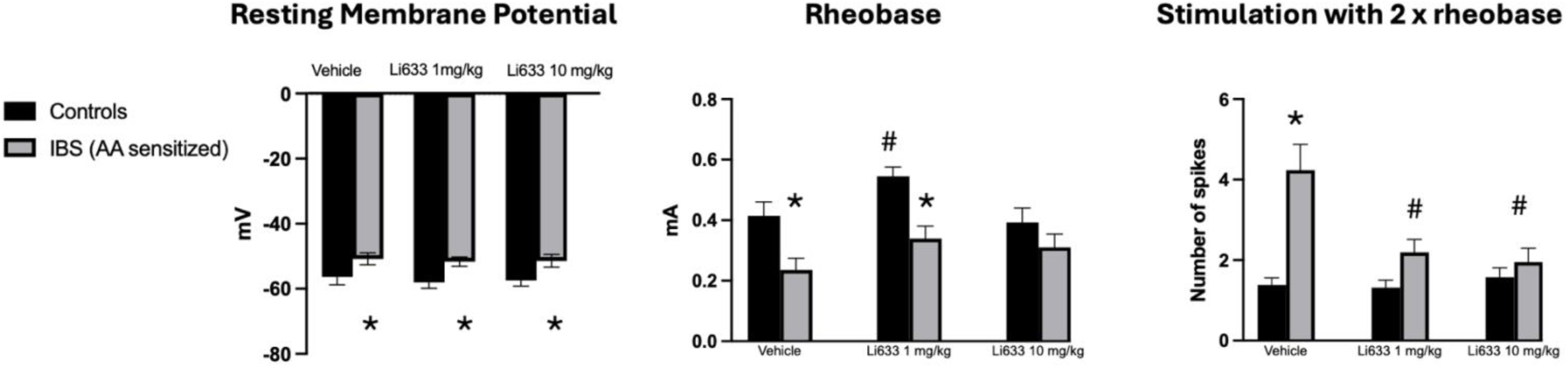
Effects of Li633 at two different doses on rats sensitized with AA in the neonatal period to produce an IBS-like model of colonic hyperalgesia. Colonic sensory neurons labeled with DiI (see methods) were dissociated from lumbosacral DRGs and excitability studied by patch clamping. **(a)** Resting membrane potential; **(b)** Rheobase (minimum amount of current to elicit an action potential spike); and **(c)** Number of spikes after current stimulation with 2x rheobase. Data are presented as mean ± SEM (n=10-13). *: Significantly different from saline group with same treatment. #: Significantly different from AA/vehicle group, P < 0.05 by Student Newman-Keuls test.

### Visceromotor response to noxious gastric distension in a rat model of FD

To further test whether LI633 also reduces hyperalgesia induced by other disorders, we tested the effect of acute treatment with LI633 in a rat model of FD.^20^ Sensitized and control rats were treated with LI633 (10 mg/kg, PO). One hour after administration, visceral hypersensitivity of the rats was assessed by the VMR responses to gastric distention at pressures of 20, 40, 60 and 80 mmHg. As shown in Figure 5, VMR responses were significantly increased in FD rats but treatment with LI633 significantly attenuated this. Two-way ANOVA revealed the main effect of treatment: F_(3,91)_ = 25.048, P< 0.001; main effect of pressure: F_(3,91)_ = 51.045, P< 0.001; Effect of treatment x pressure: F_(9,91)_ = 3.086, P = 0.003. Post hoc test (Student Newman-Keuls test) showed that FD-Vehicle group was significantly different from control-vehicle group (P < 0.05), whereas FD-LI633 group was significantly different from FD-vehicle groups (P < 0.05) but no difference from control-vehicle group.

**Figure 5.**
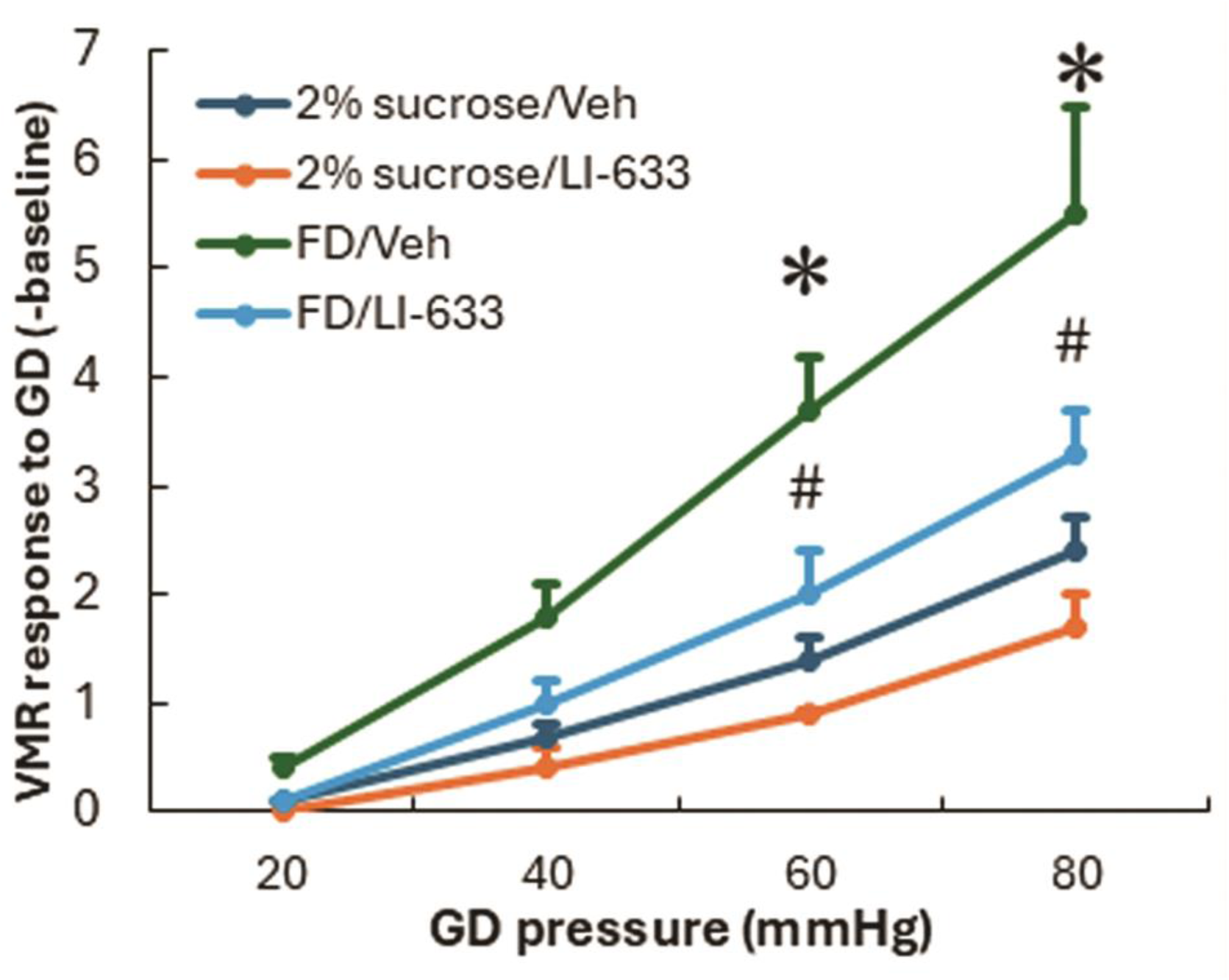
Acute treatments of LI633 significantly reduced hyperalgesia in a rat model of functional dyspepsia (FD) as assessed by visceral motor reflex (VMR) response to gastric extension (GD) with 20, 40, 60 and 80 mmHg pressures. Data are presented as mean ± SEM (n=6-7). *: Significantly different from saline group with same treatment. #: Significantly different from FD/vehicle group, P < 0.05 by Student Newman-Keuls test.

## Discussion

The gut-brain axis involves intricate bidirectional communication through multiple pathways, including neural, endocrine, immune, and metabolic routes. This complexity makes it difficult to isolate and study individual components or mechanisms. In particular, when dealing with disorders affecting this axis, it becomes challenging to separate the central versus peripheral effects of drugs that modulate neurotransmitters, given that several of these are present in both nervous systems. As an example, the role of GABA, although abundantly expressed in the gut along with its receptors, has received less attention in part because current pharmacological agents can reach therapeutic concentrations in both the central and peripheral nervous systems. An important step towards understanding the pathogenic role and therapeutic potential of GABAergic signaling in the periphery therefore is the development of pharmacological tools that can clearly separate the peripheral from the central effects. If peripheral GABA signaling is also shown to have a prominent and independent effect on pain, this will provide a robust foundation for developing drugs that can avoid off-target adverse effects in the CNS.^2^

Towards this end, here we have developed a peripherally restricted GABA_A_ PAM, LI-633. This compound binds specifically to the GABA_A_ receptor, with nanomolar affinity to its expected benzodiazepine site, in the classical flunitrazepam displacement binding assay. Further functional electrophysiological characterization using multiple overexpressing cell lines confirmed that LI-633 is devoid of direct GABA agonist activity at all concentrations tested but showed potent potentiation of GABA-evoked current, consistent with being a positive allosteric modulator (PAM) of GABA_A_. Unsurprisingly, the “benzodiazepine-insensitive” α_4_ and α_6_ containing receptors^25^ showed no potentiation in the presence of LI-633 (data not shown).

Across the various receptor subunit combinations that were examined, LI-633 demonstrated little to no selectivity to individual GABA_A_ receptor subunits, similar to most benzodiazepines. The maximum current amplitude evoked in the presence of LI-633 is comparable to that produced by diazepam across all subtypes tested, which is frequently used as a reference point for comparison. Additional screening against a diverse panel of pharmacological targets revealed no major off-target effects by LI-633.

In vitro ADME profiling revealed that LI-633 has good metabolic stability in both rat and human microsomes, relatively high unbound fraction in plasma, and no measurable inhibition of 5 cytochrome P450 enzymes, suggesting broad utility for in vivo testing. In MDR1-MDCK transfected cells, LI-633 showed low passive permeability, as well as moderate efflux by the transporter P-gp. It has been suggested that small-molecule drugs possessing this combination of properties have a lower likelihood of crossing the blood-brain barrier.^26^ Indeed, the unbound brain to plasma ratio (K_puu_) after even a very high oral dose of 100 mg/kg shows that only ∼2% of circulating drug reaches the CNS. Furthermore, LI-633 did not depress locomotor activity following doses up to 30 mg/kg, suggesting that CNS effects would not confound further efficacy experiments.

The direct ability of LI-633 to modulate GABAergic signaling in the periphery was broadly evaluated using native, labeled and isolated, rat DRG sensory neurons, which express a diverse array of GABA_A_ subunits. The EC_50_ obtained in rat DRGs (Figure 2A) was in broad agreement with the data obtained by electrophysiology in cell lines. However, an over-reliance on rodent models has been implicated in the poor success rate in translating new drug candidates to the clinic^27^, and this concern prompted us to examine the effects of LI-633 in an ex-vivo preparation of human DRG neurons. These experiments demonstrated that both muscimol and GABA were capable of reducing DRG neuron excitability when stimulated with EFS. Likewise, LI-633 potentiated the effect of submaximal GABA in a dose-dependent manner. This series of experiments confirms not only that functional GABA_A_ receptors are present on human DRG neurons, but also that by activating these receptors, their action potential firing is reduced.

Our results are in good agreement with the findings of Du et al^12^, who showed that GABA application to rat DRG neurons generally decreased the number of action potentials fired under current-clamp conditions. Interestingly, these authors also showed that GABA depolarizes most DRG neurons. Although seemingly paradoxical, reduced excitability stemming from depolarization of sensory neurons has been postulated to underlie the congenital insensitivity to pain experienced by carriers of sodium channel Na_v_1.9 gain-of-function mutations.^28^ In these individuals, persistent activation of a sodium channel leads to a large depolarization of resting membrane potential, which in turn inactivates a significant population of other sodium channels, leading to the failure of action potentials to propagate. It is tempting to speculate that a similar mechanism may be operative in the case of GABA activation, but more work is required to clarify the mechanism of action. Regardless of mechanism, the net effect of GABA_A_ activation is to reduce excitability. To our knowledge, this is the first demonstration that the excitability of human DRG neurons is reduced by GABA_A_ activation.

Proof of concept studies were then carried out to demonstrate the translation of these in vitro effects to in vivo analgesic efficacy. We chose IBS as a potential target because of several reasons. First, IBS is a very prevalent gastrointestinal disorder affecting up to 6% of the US population, significantly impacting quality of life and imposing a substantial socio-economic burden.^29^ Humans with IBS have reduced levels of glutamic acid decarboxylase (GAD), the enzyme that is responsible for GABA production along with reduced GABA levels in their colons.^30^ Secondly, several benzodiazepines have been shown to be effective in relieving so-called functional gastrointestinal symptoms including pain, ^31, 32^ and one benzodiazepine (in combination with an antispasmodic) containing formulation (chlordiazepoxide/clidinium) is approved by the FDA for the treatment of irritable bowel syndrome (IBS), a condition characterized by abdominal pain and altered bowel movements.^33^ Human experience is backed by preclinical studies which show that the GABA-A agonist muscimol reduces both colonic afferent excitability in response to stretch as well as behavioral pain responses to colonic distention.^13^

Further, both diazepam and GABA attenuate visceral hypersensitivity in mice with acute colitis,^14^ although opposite results have also been reported^34^ probably due to differences in DSS dose and route of delivery of GABA agonists (oral versus intraperitoneal). However, acute inflammation is not a feature of IBS. We therefore used a model that reliably produces hypersensitivity to colorectal distension, even in the absence of overt tissue injury or inflammation.^16, 35^ Sensitized animals showed significantly reduced behavioral responses to colorectal distention after treatment with LI-633 at oral doses as low as 1 mg/kg. From the pharmacokinetic data presented in Figure 1C, this dose is expected to produce unbound plasma concentrations of 52 nM at 1-hour post-dose. Thus, a plasma concentration at, or slightly below, the rat DRG EC_50_ is sufficient for analgesic efficacy in this model. Examination of the buprenorphine-treated group shows a nearly complete loss of sensitivity to mechanical stimulus indicating an expected generalized “numbing” effect. By contrast, LI-633 reduces the response only to the level of unsensitized animals. This suggests that LI-633 may reverse the hypersensitivity associated with IBS, while preserving normal sensory function. A similar effect on visceral hypersensitivity was also seen in rats modeled to have functional dyspepsia (FD) which is equally, if not more, prevalent than IBS.

The finding that LI-633 exerts an antinociceptive effect after repeat dosing, even after a 24-hour washout period, can perhaps be explained by the sustained, high concentrations of drug that were observed within the intestinal tissue. A compound with low solubility and very low passive permeability is expected to create a “reservoir” of drug that is slowly absorbed over many hours, and it is likely that therapeutic concentrations are maintained within the gut even after plasma drug levels have returned to baseline.

The remarkable property of LI-633 to accumulate in colon even after plasma levels have receded, suggesting that a critical site of action is in the tissue. The absent/poor brain penetration of LI-633 indicates that antinociceptive efficacy in the IBS rat model may be a result of targeting peripheral sensory receptors located within the colon; however, experiments to date cannot conclusively establish an additional role at the DRG cell body, which is likely as suggested by the study by Du et al using somatic pain models.^12^ Our results also demonstrate decreases excitability of the colon specific sensory neuronal bodies after in vivo treatment with LI633. Regardless of whether GABA_A_ signaling attenuates nociception at the cell body or at the axon (or both), this study has established that a peripheral mechanism is important in enhancing signals to the brain in a model of visceral hypersensitivity.

Although this study focused on the effects of peripheral GABA_A_ signaling on nociception, GABA likely plays an important role in gastrointestinal motility as well with well-established mechanisms and often opposing effects in both the central and enteric nervous systems.^36, 37^ With the use of tools such as LI-633 it is anticipated that these actions can now be further clarified.

In summary, the present data demonstrate that LI-633 is a potent and selective PAM of GABA_A_ receptors, with robust reductions in the excitability of both rodent and human sensory neurons and antinociceptive activities in a rat IBS and FD model. Taken together, our data suggest that LI-633 is a much-needed tool to further understand the effects of GABAergic signaling outside the CNS. Further, these results lay the foundation for peripheral GABA signaling as an important target to modulate visceral hypersensitivity in conditions such as IBS and FD.

## Materials

### Methods

#### Rat pharmacokinetic experiments

Experiments were carried out at Pharmaron (Beijing, China). Male Sprague-Dawley rats weighing between 155 and 164 g were housed under 12-hour light/ 12-hour dark cycles and were given ad libitum access to food and water. Following a 12-hour fasting period, animals received oral doses of LI-633 (1, 3, 10, or 100 mg/kg), formulated in 5% DMSO/95% pH 8.0 phosphate-buffered saline. Serial blood sampling was carried out at 0.25, 0.5, 1, 2, 4-, 8-, 12-and 24-hours post-dose. After centrifuging to obtain plasma, citric acid solution (440 mg/mL) was added at a rate of 10% by volume in order to stabilize acyl glucuronide metabolites, and the plasma samples were stored frozen until the time of analysis. LI-633 concentrations were determined at each time point by LC-MS/MS.

#### Nonspecific GABA Binding Assay

Tritiated flunitrazepam displacement from rat brain tissue was performed as described by Speth et al ^15^ at CEREP, Celle l’Evescault, France.

#### Manual patch-clamp in recombinant cell lines

Experiments were carried out by B’Sys GmbH (Witterswil, Switzerland). CHO cells stably expressing human GABA_A_ α_1_ß_2_γ_2_ receptors and LTK cells stably expressing human GABA_A_ α_2_ß_2_γ_2_, α_3_ß_2_γ_2_, or α_5_ß_2_γ_2_ receptors were used for all experiments. The 35 mm culture dishes upon which cells were seeded at a density allowing single cells to be recorded, were placed on the dish holder of the microscope and continuously perfused (at approximately 1 mL/min) with bath solution (137 mM Sodium Chloride, 4 mM Potassium Chloride, 1.8 mM Calcium Chloride, 1 mM Magnesium Chloride, 10 mM HEPES, 10 mM D-Glucose pH (NaOH) 7.4). All solutions applied to cells, including the pipette solution (130 mM Potassium Chloride, 1 mM Magnesium Chloride, 5 mM Mg-ATP, 10 mM HEPES, 5 mM EGTA, pH (KOH) 7.2) were maintained at room temperature (19°C to 30°C). After formation of a Gigaohm seal between the patch electrodes and an individual cell (pipette resistance range: 2.5 MΩ to 6.0 MΩ; seal resistance range: >1 GΩ) the cell membrane across the pipette tip was ruptured to assure electrical access to the cell interior (whole-cell patch-clamp configuration). GABAAR inward currents were measured upon application of submaximal GABA concentration to patch-clamped cells. The cells were voltage-clamped at a holding potential of -80 mV. If current density was judged to be too low for measurement, another cell was recorded. Only data from cells treated with the test item were documented. All GABAAR subtypes were stimulated by GABA (2 µM) and cumulative, increasing concentrations of LI-633/GABA. Between two GABA applications bath solution (without or with LI-633) was perfused for at least 60 s. For control experiments, the test item was replaced by 0.1% DMSO and a single concentration of GABA was applied at the end of the experiment. Only cells with initial current amplitudes between 150 pA and 1000 pA were used for analysis.

#### Off-target binding panel

LI-633 was tested against a panel of 87 enzymes, receptors, and ion channels (Eurofins Panlabs, Taipei, Taiwan) using methods described by Eurofine Panlabs https://www.eurofinsdiscovery.com/catalog/safetyscreen87-panel-tw/PP223. LI-633 was tested in duplicate at a concentration of 10 uM.

#### Rat dorsal root ganglia (DRG) cell electrophysiology

Experiments were carried out at Metrion Biosciences (Cambridge, UK). Dorsal root ganglion neurons were isolated from P8-P12 Sprague-Dawley rats, were prepared and maintained in culture for up to four days on glass coverslips. Whole-cell manual patch clamp recordings were obtained from cell bodies while maintaining a holding potential of -60 mV. A VC38 pressurized perfusion system (ALA Scientific Instruments) was used for solution exchange. HEKA Fitmaster, Prism, and Excel were used for data analysis. Intracellular recording solution contained the following: HEPES (10 mM), MgCl2 (2 mM), EGTA (10 mM), Mg-ATP (3 mM), CsCl (130 mM), pH adjusted to 7.2 by addition of cesium hydroxide. Extracellular recording solution contained the following: NaCl (140 mM), KCl (2.5 mM), HEPES (10 mM), MgCl_2_ (1 mM), CaCl_2_ (2 mM), glucose (10 mM), pH adjusted to 7.4 by addition of cesium hydroxide. Drugs (muscimol and LI-633) were applied for 20 seconds in order to ensure that maximum available current was activated.

#### Human DRG Electrical Field Stimulation Study

Experiments were carried out at AnaBios (San Diego, USA). Tissue preparation: Human DRGs obtained from consenting donors were transferred into a dissection vessel containing a cold (4°C), fresh proprietary dissection solution. DRGs were maintained completely submerged in dissection solution and dissected appropriately. Each DRG was enzymatically dissociated as per AnaBios’ proprietary methodologies. Dissociated cells were seeded on 96-well plastic bottom plates (Corning) that had been precoated with poly-D-lysine. Cells were maintained in culture at 37°C with 5% CO2 in 200 µL DMEM/F12 supplemented with 10% horse serum (Thermo Fisher Scientific), 2 mM glutamine, 25 ng/mL hNGF (Cell Signaling Technology), 25 ng/mL GDNF (Peprotech), Gem21 NeuroPlex (GeminiBio) and penicillin/ streptomycin (Thermo Fisher Scientific). Half of the culture media was replaced with fresh media every 3 days.

Recording Procedure: The calcium dye, 3uM Fluo-8 AM (AAT Bioquest) in calcium imaging buffer was loaded in the well for a period of 20 to 25 min. Fluo-8-loaded cells were excited at 480 nm and emission was collected at 520 nm with a pcoEDGE sCMOS camera (PCO) mounted on an inverted microscope (Olympus IX71). The temperature of the solution was maintained at room temperature. The DRG neurons were tested in optical electric field stimulation (EFS) recording using a stimulator (Master 9 AMPI). The baseline excitability profile of the cells was first assessed with a train of 10 stimuli delivered at a voltage between 1500-2000 mV at 5Hz. Following the baseline profiling, the cells were exposed to the vehicle or LI-633 by perfusion at 1 mL/min for 2 minutes, followed by EFS with 2 second recording. Then, the cells were washed out for 5 min and repeated the above procedure with various concentration of LI-633 and vehicle. At the end of the of the EFS recordings, a positive control, 200 nM capsaicin, was applied by perfusion at 1 mL/min for 3 min to establish the cells’ sensitivity to that common nociceptive agent. Recordings were performed in streaming mode at 50Hz for the EFS 5Hz part- and in-time lapse mode at 0.2Hz for the capsaicin application. For each condition, the number of cells activated versus the appropriate control was counted.

Cell exclusion criteria: Absence of calcium response following stimulation, time frame of drug exposure not respected, unstable cell response (i.e., baseline exhibited a decrease over 10% during a 10-pulse stimulation), or lack of response to EFS.

## Author Contributions

MSP: Investigation; supervision; visualization; writing of original draft

QL: Investigation; supervision; formal analysis; visualization

IPB: Project administration; writing (review and editing)

YF: Investigation

LL: Investigation

YZ: Investigation; formal analysis

SK: Investigation; formal analysis

GC: Formal analysis; writing (review and editing)

AD: Investigation

NW: Investigation

DP: Writing (review and editing)

JCB: Methodology; supervision; formal analysis; drug design, writing of original draft; writing (review and editing); resources; funding acquisition

PJP: Conceptualization; methodology; supervision; formal analysis; drug design, writing of original draft; writing (review and editing); resources; funding acquisition

---

**Table S1:** In vitro ADME properties of LI-633.

|  |  |
| --- | --- |
| Kinetic sol. pH 7.4 ($\mu\text{g/mL}$) | 125 |
| MDR1-MDCK $P_{\text{app}}$ A-B (nm/s) | 3.1 |
| MDR1-MDCK Efflux Ratio | 2.8 |
| R,H hepatocytes $T_{1/2}$ (min) | >120 |
| R,H microsomes $T_{1/2}$ (min) | >120 |
| Rat PPB, % unbound | 18.6 |
| Human PPB, % unbound | 7.5 |
| Rat BTB, % unbound | 13.1 |
| CYP450 $\text{IC}_{50}$, $\mu\text{M}$ | >50 |

**Figure S1.**
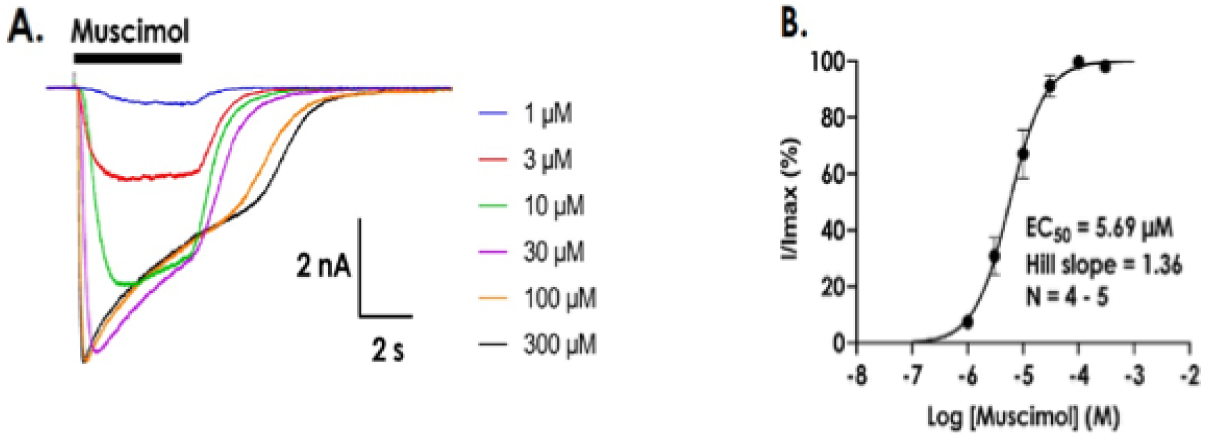
GABAergic currents elicited by muscimol using whole cell manual patch clamp recordings were made DRG cell bodies. **(A)** Representative currents shown with 4 second agonist exposure. Maximal current activation demonstrated with 100 µM (no further increase with 300 µM). Currents showed expected activation & inactivation characteristics. Activation was more rapid with higher concentrations and desensitization increased as with higher concentrations of muscimol. (B) Mean I/Imax data (±S.D) with fitted concentration response curve.

## References

1. Basbaum AI, Bautista DM, Scherrer G, Julius D. Cellular and molecular mechanisms of pain. Cell 139, 267–284 (2009).

2. Apkarian AV, Reckziegel D. Peripheral and central viewpoints of chronic pain, and translational implications. Neurosci Lett 702, 3–5 (2019).

3. Sperber AD, et al. Worldwide Prevalence and Burden of Functional Gastrointestinal Disorders, Results of Rome Foundation Global Study. Gastroenterology 160, 99–114 e113 (2021).

4. Mittal R, et al. Neurotransmitters: The Critical Modulators Regulating Gut-Brain Axis. J Cell Physiol 232, 2359–2372 (2017).

5. Sieghart W, Savić MM. International Union of Basic and Clinical Pharmacology. CVI: GABAA receptor subtype-and function-selective ligands: key issues in translation to humans. Pharmacological reviews 70, 836–878 (2018).

6. Sieghart W. Structure, Pharmacology, and Function of GABA(A) Receptor Subtypes. Adv Pharmacol 54, 231–263 (2006).

7. Miki Y, Taniyama K, Tanaka C, Tobe T. GABA, glutamic acid decarboxylase, and GABA transaminase levels in the myenteric plexus in the intestine of humans and other mammals. J Neurochem 40, 861–865 (1983).

8. Auteri M, Zizzo MG, Serio R. GABA and GABA receptors in the gastrointestinal tract: from motility to inflammation. Pharmacol Res 93, 11–21 (2015).

9. McCarson KE, Enna S. GABA pharmacology: the search for analgesics. Neurochemical research 39, 1948–1963 (2014).

10. Zeilhofer HU, Ralvenius WT, Acuna MA. Restoring the spinal pain gate: GABA(A) receptors as targets for novel analgesics. Adv Pharmacol 73, 71–96 (2015).

11. Knabl J, et al. Reversal of pathological pain through specific spinal GABAA receptor subtypes. Nature 451, 330–334 (2008).

12. Du X, et al. Local GABAergic signaling within sensory ganglia controls peripheral nociceptive transmission. The Journal of clinical investigation 127, 1741–1756 (2017).

13. Loeza-Alcocer E, McPherson TP, Gold MS. Peripheral GABA receptors regulate colonic afferent excitability and visceral nociception. J Physiol 597, 3425–3439 (2019).

14. Gold MS, Loeza-Alcocer E. Experimental colitis-induced visceral hypersensitivity is attenuated by GABA treatment in mice. Am J Physiol Gastrointest Liver Physiol 326, G252–G263 (2024).

15. Speth RC, Wastek GJ, Yamamura HI. Benzodiazepine receptors: temperature dependence of [3H] flunitrazepam binding. Life sciences 24, 351–357 (1979).

16. Winston J, Shenoy M, Medley D, Naniwadekar A, Pasricha PJ. The vanilloid receptor initiates and maintains colonic hypersensitivity induced by neonatal colon irritation in rats. Gastroenterology 132, 615–627 (2007).

17. Xu GY, Winston JH, Shenoy M, Zhou S, Chen JD, Pasricha PJ. The endogenous hydrogen sulfide producing enzyme cystathionine-beta synthase contributes to visceral hypersensitivity in a rat model of irritable bowel syndrome. Mol Pain 5, 44 (2009).

18. Zhu Y, et al. Nerve growth factor modulates TRPV1 expression and function and mediates pain in chronic pancreatitis. Gastroenterology 141, 370–377 (2011).

19. Cordner ZA, et al. Vagal gut-brain signaling mediates amygdaloid plasticity, affect, and pain in a functional dyspepsia model. JCI Insight 6, (2021).

20. Liu LS, Winston JH, Shenoy MM, Song GQ, Chen JD, Pasricha PJ. A rat model of chronic gastric sensorimotor dysfunction resulting from transient neonatal gastric irritation. Gastroenterology 134, 2070–2079 (2008).

21. Masiulis S, et al. GABA(A) receptor signalling mechanisms revealed by structural pharmacology. Nature 565, 454–459 (2019).

22. Hollands EC, et al. Population patch-clamp electrophysiology analysis of recombinant GABAA alpha1beta3gamma2 channels expressed in HEK-293 cells. J Biomol Screen 14, 769–780 (2009).

23. Ebert B, Thompson SA, Saounatsou K, McKernan R, Krogsgaard-Larsen P, Wafford KA. Differences in agonist/antagonist binding affinity and receptor transduction using recombinant human gamma-aminobutyric acid type A receptors. Mol Pharmacol 52, 1150–1156 (1997).

24. Fouillet A, et al. Characterisation of Nav1. 7 functional expression in rat dorsal root ganglia neurons by using an electrical field stimulation assay. Molecular Pain 13, 1744806917745179 (2017).

25. Maramai S, Benchekroun M, Ward SE, Atack JR. Subtype selective γ-aminobutyric acid type A receptor (GABAAR) modulators acting at the benzodiazepine binding site: an update. Journal of Medicinal Chemistry 63, 3425–3446 (2019).

26. Bagal SK, Bungay PJ. Minimizing Drug Exposure in the CNS while Maintaining Good Oral Absorption. Acs Med Chem Lett 3, 948–950 (2012).

27. Yekkirala AS, Roberson DP, Bean BP, Woolf CJ. Breaking barriers to novel analgesic drug development. Nat Rev Drug Discov 16, 544–563 (2017).

28. Huang J, et al. Sodium channel Na V 1.9 mutations associated with insensitivity to pain dampen neuronal excitability. The Journal of clinical investigation 127, 2805–2814 (2017).

29. Almario CV, Sharabi E, Chey WD, Lauzon M, Higgins CS, Spiegel BMR. Prevalence and Burden of Illness of Rome IV Irritable Bowel Syndrome in the United States: Results From a Nationwide Cross-Sectional Study. Gastroenterology 165, 1475–1487 (2023).

30. Aggarwal S, Ahuja V, Paul J. Dysregulation of GABAergic Signalling Contributes in the Pathogenesis of Diarrhea-predominant Irritable Bowel Syndrome. J Neurogastroenterol Motil 24, 422–430 (2018).

31. Baume P, Cuthbert J. The effect of medazepam in relieving symptoms of functional gastrointestinal distress. Aust N Z J Med 3, 457–460 (1973).

32. Baume P, Tracey M, Dawson L. Efficacy of two minor tranquilizers in relieveing symptoms of functional gastrointestinal distress. Aust N Z J Med 5, 503–506 (1975).

33. Salari P, Abdollahi M. Systematic review of modulators of benzodiazepine receptors in irritable bowel syndrome: Is there hope? World J Gastroentero 17, 4251–4257 (2011).

34. Ma X, et al. Activation of GABA(A) Receptors in Colon Epithelium Exacerbates Acute Colitis. Front Immunol 9, 987 (2018).

35. Al–Chaer ED, Kawasaki M, Pasricha PJ. A new model of chronic visceral hypersensitivity in adult rats induced by colon irritation during postnatal development. Gastroenterology 119, 1276–1285 (2000).

36. Auteri M, Zizzo MG, Serio R. GABA and GABA receptors in the gastrointestinal tract: from motility to inflammation. Pharmacological research 93, 11–21 (2015).

37. Seifi M, et al. Molecular and functional diversity of GABA-A receptors in the enteric nervous system of the mouse colon. Journal of Neuroscience 34, 10361–10378 (2014).

